# TDRD3 functions as a selective autophagy receptor with dual roles in autophagy and modulation of stress granule stability

**DOI:** 10.1101/2024.09.22.614367

**Authors:** Matthew Deater, Richard E. Lloyd

## Abstract

Tudor Domain Containing 3 (TDRD3) is a methylarginine-reader protein that functions as a scaffold in the nucleus facilitating transcription, however TDRD3 is also recruited to stress granules (SGs) during the Integrated Stress Response (ISR) although its function therein remains largely unknown. We previously showed that TDRD3 is a novel antiviral restriction factor that is cleaved by virus 2A protease, and plays complex modulatory roles in both interferon and inflammatory signaling during stress and enterovirus infections. Here we have found that TDRD3 contains structural motifs similar to known selective autophagy receptors such as p62/SQSTM1, sharing ubiquitin associated domains (UBA) and LC3 interacting regions (LIR) that anchor cargo destined for autophagosomes to activated LC3 protein coating autophagosome membranes. This is of interest since enteroviruses hijack autophagy machinery to facilitate formation of viral replication factories, virus assembly and egress from the infected cell. Here we explored possible roles of TDRD3 in autophagy, hypothesizing that TDRD3 may function as a specialized selective autophagy receptor. We found that KO of TDRD3 in HeLa cells significantly reduces starvation induced autophagy, while its reintroduction restores it in a dose-dependent manner. Autophagy receptors are degraded during autophagy and expression levels decrease during this time. We found that TDRD3 levels decrease to the same extent as the autophagy receptor p62/SQSTM1 during autophagy, indicating autophagy-targeted turnover in that role. Knockout of TDRD3 or G3BP1 did not make significant changes in overall cell localization of LC3B or p62/SQSTM1, but did result in greater concentration of Lamp2 phagosome marker for phagosomes and phagolysosomes. To test the potential roles of TDRD3 in autophagic processes, we created a series of deletion mutants of TDRD3 lacking either UBA domain or the various LIR motifs that are predicted to interact with LC3B. Microscopic examination of starved cells expressing these variants of TDRD3 showed ΔLIR-TDRD3 had defects in colocalization with LC3B or Lamp2. Further, super resolution microscopy revealed ring structures with TDRD3 interfacing with p62/SQSTM1. In examination of arsenite induced stress granules we found recruitment of TDRD3 variants disrupted normally tight SG condensation, altered the decay rate of SGs upon release from stress and the kinetics of SG formation. We found evidence that the LIR3 motif on TDRD3 is involved in TDRD3 interaction with LC3B in coIP experiments, colocalization studies, and that this motif plays a key role in TDRD3 recruitment to SGs and SG resolution. Overall, these data support a functional role of TDRD3 in selective autophagy in a mode similar to p62/SQSTM1, with specific roles in SG stability and turnover. Enterovirus cleavage of TDRD3 likely affects both antiviral and autophagic responses that the virus controls for replication.

## Introduction

TDRD3 is a multifunctional and multidomain scaffolding protein described initially as a methyl reader protein that binds asymmetric dimethylarginine marks on the RG-rich domains of multiple proteins via its TUDOR domain^1–3^. TDRD3 has an N-terminal oligonucleotide-binding (OB) fold domain and a Tudor domain that recognizes methyl arginine on target proteins. TDRD3 is expressed in most tissues and is found in both the nucleus and cytosol of the cells. In the nucleus, the Tudor domain recognizes two major types of active methyl-arginine histone marks and plays important roles in transcriptional activation, and may play roles in cancer and other diseases^3,4^. Interactions with one of its binding targets, topoisomerase 3B (TOP3B), stimulates TOP3B activity in R-loop resolution and chromatin targeting^5–7^. TDRD3 was also found to enter cytoplasmic stress granules (SG), with mostly unknown functions within that compartment(^1–3^). Recently, PRMT1 and TDRD3 were found to promote SG assembly by rebuilding protein-RNA interaction networks through PRMT1 directed methylation of SG RNA binding proteins, recruiting TDRDQ ^8^.

We have recently shown that TDRD3 is also a novel antiviral restriction factor versus enteroviruses that can promote stress granule formation and activation of a complex spectrum of interferon activated genes, alpha and lambda interferons and NFkB ^9^. We searched for mechanisms by which TDRD3 might restrict virus replication or modulate stress granule function, but could not find a role for TDRD3 interactions with G3BP1 through methylation of its RGG domain in SG assembly ^9^. Enteroviruses block the formation of stress granules through cleavage of the major SG nucleator protein G3BP1 and also cleave TDRD3 with viral 2A and 3C proteinases^9–11^. Enteroviruses and other positive strand RNA viruses also hijack autophagy machinery to facilitate formation of viral replication factories, promote RNA replication, virus assembly and egress from the infected cell^12,13^. Enteroviruses cleave the autophagy receptor p62/SQSTM1, recruit proLC3 or LC3-1 to replication organelles for replication, blocking completion of autophagosome formation^12,14,15^. Further, autophagy is known to be a mechanism to repress SG stability, and is proposed to eliminate SGs at the resolution of cell stress^16–18,19,20^.

While it has been known for some time that TDRD3 possesses a ubiquitin-associated domain (UBA)^1^ we have found, via short linear motif scanning, that it also possesses three LC3 interaction regions (LIRs), raising the possibility that TDRD3 functions in autophagy. These domains are known to be functionally required for selective autophagy receptors. Here we show that both UBA and LIR domains of TDRD3 are functional in modulating autophagy in cells, and propose that TDRD3 is a novel type of selective autophagy receptor that modulates SG assembly and possibly functions in their eventual degradation at the successful completion of the integrated stress response.

## Results

### TDRD3 Architecture Suggests that TDRD3 Functions in Autophagy

Analysis of TDRD3 structure for small linear motifs of known function (ELM)^21^ predicts that TDRD3 contains three types of key functional domains and linear amino acid motifs that are shared with archetypical selective autophagy receptors (SARS) **(Fig. 1A)**^**22**^. This includes the UBA ubiquitin-binding domain that engages ubiquitinated substrates, LC3 interacting regions (LIRs), and Traf interaction motifs. One of these LIRs (LIR #1) resides within TDRD3’s putative ssRNA/DNA binding OB-Fold domain, while the second resides within a highly disordered region (LIR #2), and the third exists flanking the upstream region of TDRD3’s methylarginine reading Tudor domain (LIR #3). This prompted us to explore if TDRD3 played any roles in autophagy processes.

**Fig. 1.**
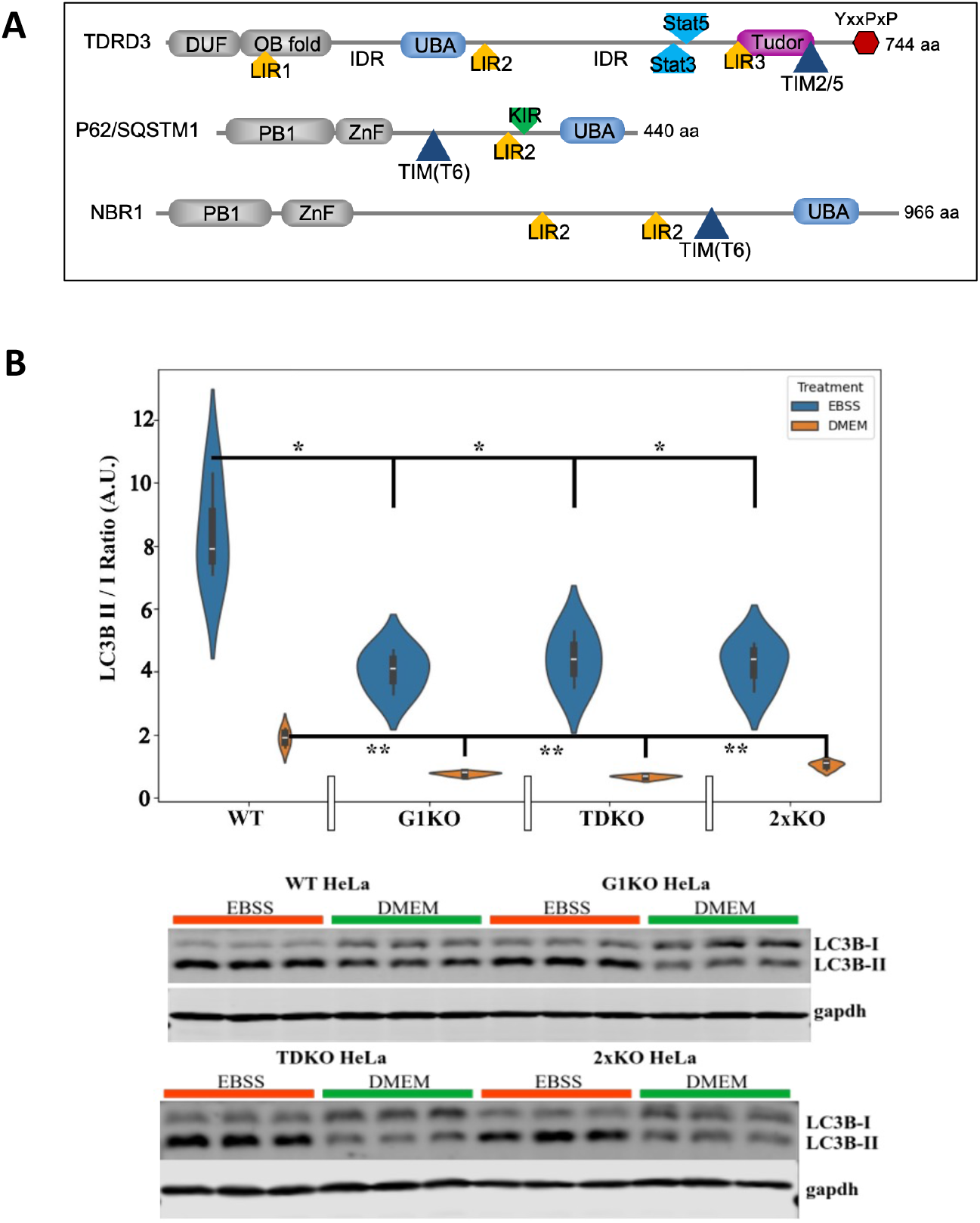
A. Domain structure of TDRD3 and archetype autophagy receptors. TDRD3 domains include DUF(domain of unknown function), OB fold, Tudor domain and two large intrinsically disordered regions (IDR). TDRD3 has three functional domains and motifs shared with key autophagy receptors p62/SQSTM1 and NBR1; LIR motif (LC3A/B/C interaction motif), UBA ubiquitin binding domain and Traf interaction motifs (TIM). LIR binds AT8 family proteins during autophagy. B. Both TDRD3 and G3BP1 facilitate basal and starvation-induced autophagy. Wild-type HeLa cells or knockouts for G3BP1 (G1KO), TDRD3 (TDKO) or both G3BP1 and TDRD3 (2xKO) were plated in triplicate, grown in DMEM supplemented with 10% FBS until confluency. Some plates were starved by washing in PBS then incubation in Earle’s Balanced Salt Solution (EBSS) for 4 hrs. All cells were collected for analysis, quickly lysed in RIPA buffer, sonicated briefly and analyzed by western blot against LC3BI and LC3BII. LC3B ratios were determined by densitometry and statistics performed using T-tests between cell types of similar treatments. Representative Western blots are shown below. *=p ≤ 0.05, **=p ≤ 0.01, ***=p ≤ 0.001.

### Both TDRD3 and G3BP1 Function in Starvation-Induced Autophagy

We first examine whether or not either TDRD3 or G3BP1 play any role in autophagy during starvation conditions. To this end, we generated HeLa cells that are knockouts for either G3BP1 (G1KO), TDRD3 (TDKO) or G3BP1^-/-^TDRD3^-/-^(2xKO) using CRISPR/Cas9 targeting either G3BP1 or all three isoforms of TDRD3. Knockouts were confirmed via WB **(Suppl. Fig1)**. We induced starvation and autophagy responses by incubation of cells with Earle’s Balanced Salt Solution (EBSS) for 4 hours or untreated (DMEM) - either of which was supplemented with 20 mM of Ammonium Chloride widely known to block autophagolysosomal degradation of cargo including LC3B-II, ultimately increasing observable LC3B-II signal via western blot. Active autophagy detection was performed by analysis of the ratio between unprocessed soluble LC3B-I and PE-conjugated membrane-bound LC3B-II ^23^.

We found that KO of either G3BP1 or TDRD3 (or both) resulted in significantly reduced active autophagy compared to WT cells, approximately two-fold. Examination of basal autophagy levels in untreated (DMEM) cells also revealed similarly reduced levels of basal autophagy as expected, evidenced by reduced LC3B-II/LC3B-1 ratios in all cases, but showing the same relative trend of reduced autophagy in KO cells **(Fig. 1B)**. Interestingly, either G3BP1 KO or TDRD3 KO reduced autophagy. Since all cell types showed increased processed LCB3-II after autophagy induction/starvation compared to their DMEM-treated counterparts, we conclude that autophagy in these cells is not completely restricted but instead impaired. Overall, these data suggest that in the absence of either TDRD3 or G3BP1 (or both), autophagy is significantly inhibited compared to WT cells and that this effect is present during both starvation (possibly representing nonselective microautophagy) and basal autophagy conditions (possibly representing selective autophagy).

### TDRD3 and the SAR p62/SQSTM1 display coordinate expression during autophagy initiation and inhibition

Next we wanted to see if TDRD3 and p62/SQSTM1 follow similar expression patterns during autophagy as p62/SQSTM1, in accordance with its function as a selective autophagy receptor. p62/SQSTM1 is rapidly degraded during autophagy its total expression level is reduced, thus it can serve as a measure of autophagic activity^24^. To do this, we starved WT HeLa cells with EBSS supplemented with ammonium chloride as previously described. Both TDRD3 and p62/SQSTM1 expression levels markedly decreased during starvation compared to DMEM treated cells (DMEM only vs EBSS only)**(Fig. 2A)**. Additionally, the expression of both TDRD3 and p62/SQSTM1 increased when basal autophagy is inhibited (DMEM + NH_4_Cl), and both are protected to nearly WT levels when protected from proteasome activity (MG132). This suggests that both TDRD3 and p62/SQSTM1 are being degraded during autophagy possibly due to LC3 interactions on the autophagosome membrane, ubiquitin interactions and subsequent degradation along with their presumptive cargo. It is important to note that TDRD3 is known to be a transcriptional coactivator through binding to methyl-arginine residues in histones ^2,25^). Thus we tested if reintroduction of TDRD3 altered transcription levels of p62/SQSTM1 or LC3B by RT-qPCR assay. Results indicated no significant alteration of p62/SQSTM1 or LC3B RNA levels in cells were detected (Supplemental Fig. 2).

**Fig. 2.**
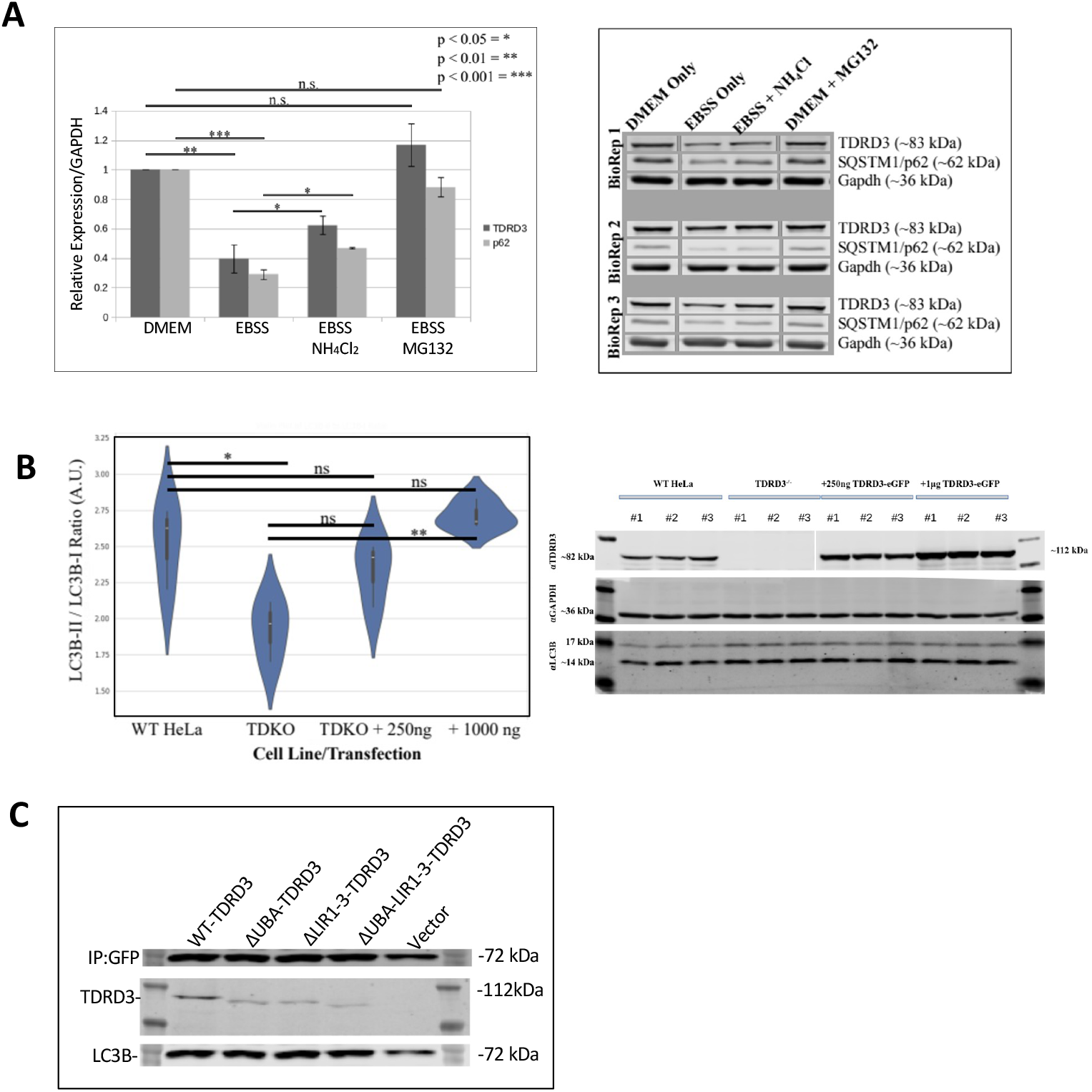
A. TDRD3 and p62/SQSTM1 display coordinate altered expression levels during autophagy induction and inhibition. Cells were untreated (DMEM), or starved by washing in PBS then incubation in Earle’s Balances Salt Solution (EBSS) for 4 hrs. Alternatively, cells starved with EBSS supplemented with either 20 mM NH_4_Cl or 50 μg MG132 per manufacturer’s instruction. to inhibit autophagic protein turnover (autophagic flux). Cells were analyzed by western blot for LC3BI/LC3BII. LC3B ratios were determined by densitometry and statistics performed using T-tests between cell types of similar treatments. *=p ≤ 0.05, **=p ≤ 0.01, ***=p ≤ 0.001. B. TDRD3 rescues autophagy during starvation. WT HeLa, TDRD3 KO cells or TDRD3 KO cells ectopically expressing TDRD3-eGFP introduced at indicated plasmid levels/well were starved in EBSS for 2 hrs with no inhibition (N=3). Cells were analyzed by Western blot using antibodies for TDRD3, GAPDH or LC3B. Experiment was repeated (n=6). LC3B-II to LC3B-I ratios were calculated as a measure of autophagic potential for each condition. C. TDRD3 co-immunoprecipitates with LC3B. TDKO HeLa cells were cotransfected at a 1:1 ratio with TDRD3 variants expression plasmids lacking GFP tags along with an LC3B-GFP-RFP fusion construct for 18hr. Cells were starved with EBSS for 4 hrs and coIP was performed using Chromotec GFP-Trap and precipitates analysed by Western blot.

### Reintroduction of TDRD3 Rescues Starvation-Induced Autophagy

Next we wanted to see if reintroduction of TDRD3 could restore autophagic activity in cells during nutrient starvation. To do this, we first determined levels of TDRD3 transfection that resulted in near endogenous expression levels which was approximately 250ng/well of a 12-well plate (data not shown). Following this, we plated WT HeLa cells, TDRD3 KO HeLa cells, and TDRD3 KO HeLa cells transfected with either 250ng/well or 1000ng/well of a 12-well plate, in triplicate, twice, weeks apart. Cells were then starved for 2 hours in EBSS pH 7. Cells were then washed, lysed, and western blot against LC3B was performed. Reintroduction of TDRD3 was found to rescue autophagy in a dose-dependent manner **(Fig. 2B)**.

### TDRD3 interacts with LC3B

Since we found that TDRD3 was playing important roles in autophagy we sought evidence that TDRD3 interacted with LC3B, which has not been reported previously. We expressed a tagged version of LC3B together with inducible constructs expressing WT-TDRD3 or variants lacking the UBA(ΔUBA-TDRD3), LIR motifs1,2,3 (ΔLIR1-3-TDRD3) or both (or vector control) and performed performed GFP-trap co-immunoprecipitations against LC3B (**Fig. 2C)**. These experiments confirmed that TDRD3 interacts in a complex with LC3B. Interestingly, a loss of TDRD3 co-immunoprecipitation signal was observed with either ΔUBA or ΔLIR1-3 variants and the TDRD3 variant lacking both was recovered mostly weakly. This indicates that TDRD3 likely does interact with LC3B via its LIR motifs, that both motifs play a role in interactions between TDRD3 and LC3B in cells, although neither is solely required hinting at a possible complex of TDRD3, LC3B, and as of yet unknown interactors.

### TDRD3 and G3BP1 KO cells display altered phagosome morphology

We next compared the three TDRD3 or G3BP1 KO cell lines to WT cells using microscopic analysis of the distribution of autophagy and phagosome markers LC3B, p62/SQSTM1, and Lamp2 during either arsenite stress or starvation induced autophagy. Starvation resulted in formation of intracellular and extracellular vesicles containing both p62/SQSTM1 and LC3B **(Fig. 3A)**, however arsenite treatment resulted in only subtle changes, with modest organization of p62/SQSTM1 in larger foci **(Fig. 3B)**. Importantly, there was no significant differences observed between the KO cells and WT cells in the distribution of these markers.

**Fig. 3.**
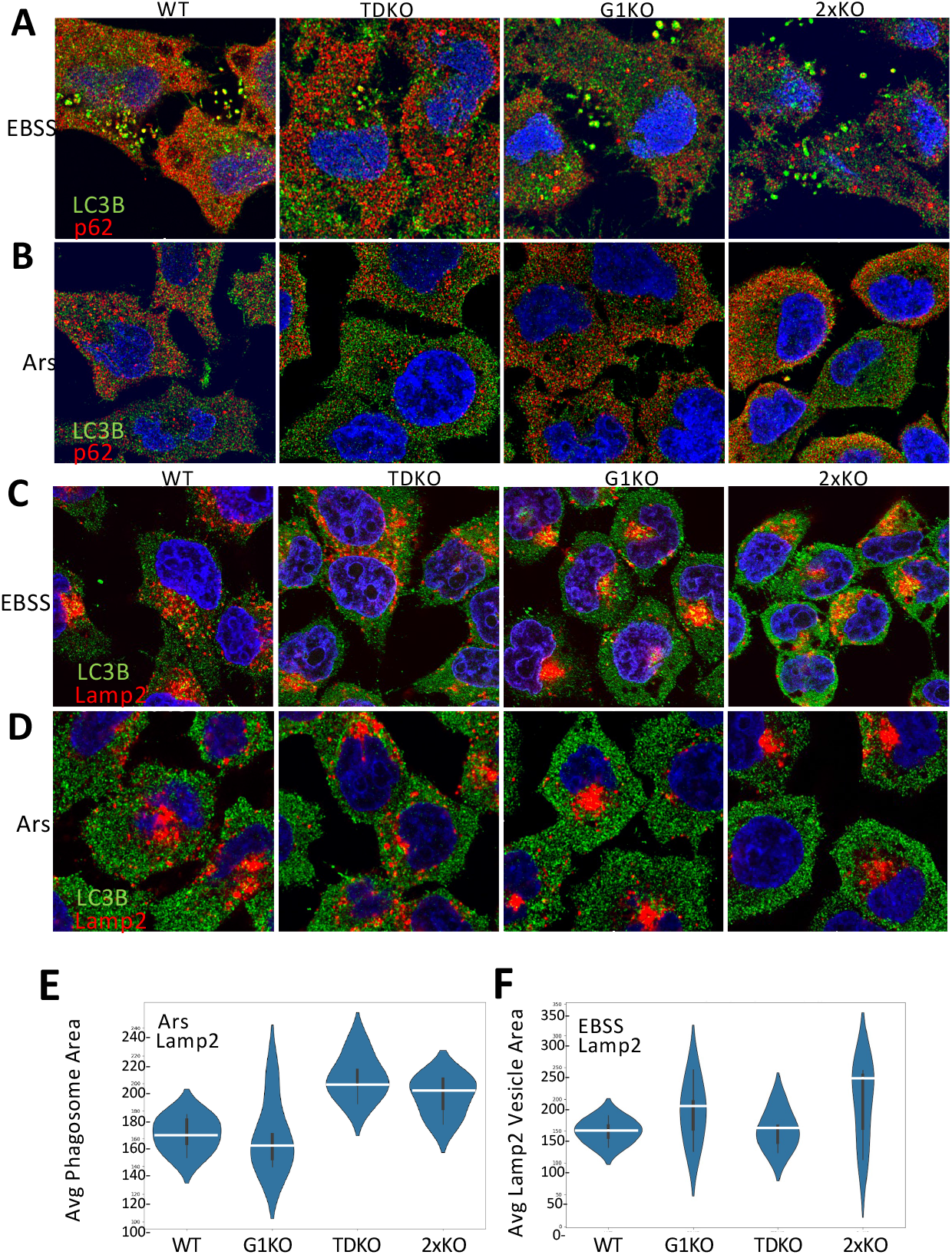
Altered morphology of LC3B/p62/SQSTM1 autophagosome and Lamp2 phagolysosomes in TDRD3 or G3BP1 knockout cells. Cells were starved 4hrs with EBSS (A, C) or treated with arsenite for 1 hr (B, D) before processing for immunofluorescence with indicated antibodies. E. Quantitation of phagosome sizes during arsenite stress. F. Quantitation of phagosome size during starvation.

Examination of lysosomes or autophagolysosomes using Lamp2 as a marker revealed differences between the cell types during both starvation and arsenite treatment. In WT cells, Lamp2 congregated on one side of the cell, and tended to concentrate into peri-nuclear degradation centers in the cytoplasm during starvation**(Fig. 3C)**. This also included LC3B, which colocalized with many Lamp2 foci. Interestingly, the distribution of foci was not as compact or condensed in TDKO cells. Conversely, cells with G3BP1 KO (GIKO and 2xKO) displayed tighter condensation than WT cells. Similar differential condensation of Lamp2 was also observed in cells treated with arsenite**(Fig. 3D)**. We examined the average size of Lamp2 phagosomes and found that during arsenite treatment TDKO and 2xKO cells displayed larger vesicles**(Fig. 3E)**. Conversely, during starvation cells with G3BP1 KO tend to make larger autophagolysosomes or lysosomes **(Fig. 3F)**. This suggests that TDRD3 may play a role in condensing or maturation of autophagolysosomes or lysosomes during autophagy, whereas, G3BP1 may play a role in inhibiting such condensation. Conversely, G3BP1 may play a role in autophagolysosome or lysosome condensation during arsenite stress.

### TDRD3 deletion variants with altered interactions with autophagosome machinery

To test if specific motifs in TDRD3 played roles during autophagy, we created a series of deletion mutant variants of TDRD3 that lacked the UBA motif, or respective LIR motifs, either singly or in combinations. When TDRD3 was expressed in TDKO cells then starved with EBSS, no significant changes were noted in LC3B and p62/SQSTM1 distribution compared to cells not expressing TDRD3, similar to results in Fig 3A. Overall, only limited TDRD3 was detected colocalizing with either p62/SQSTM1 or LC3B, particularly in autophagosomes labeled with both p62 and LC3B **(Fig. 4A)**, unless higher level expression of TDRD3 occurred. In this case, foci may represent small nascent SGs. However, various sized TDRD3 foci under lower expression conditions could be observed adjacent or partly colocalizing with p62/SQSTM1 foci. Comparison of WT-TDRD3 foci with those formed with ΔUBA or ΔLIR3 variants under starvation conditions were similar, where TDRD3 foci were closely adjacent to p62/SQSTM1/LC3B autophagosomes or LC3B foci. A subset of cells displayed larger structure that colocalized with both LC3B and p62/SQSTM1 **(Fig. 4B)** These data indicate a portion of TDRD3 may be involved in autophagosome processes, and TDRD3 variants display partly altered phenotypes.

**Fig. 4.**
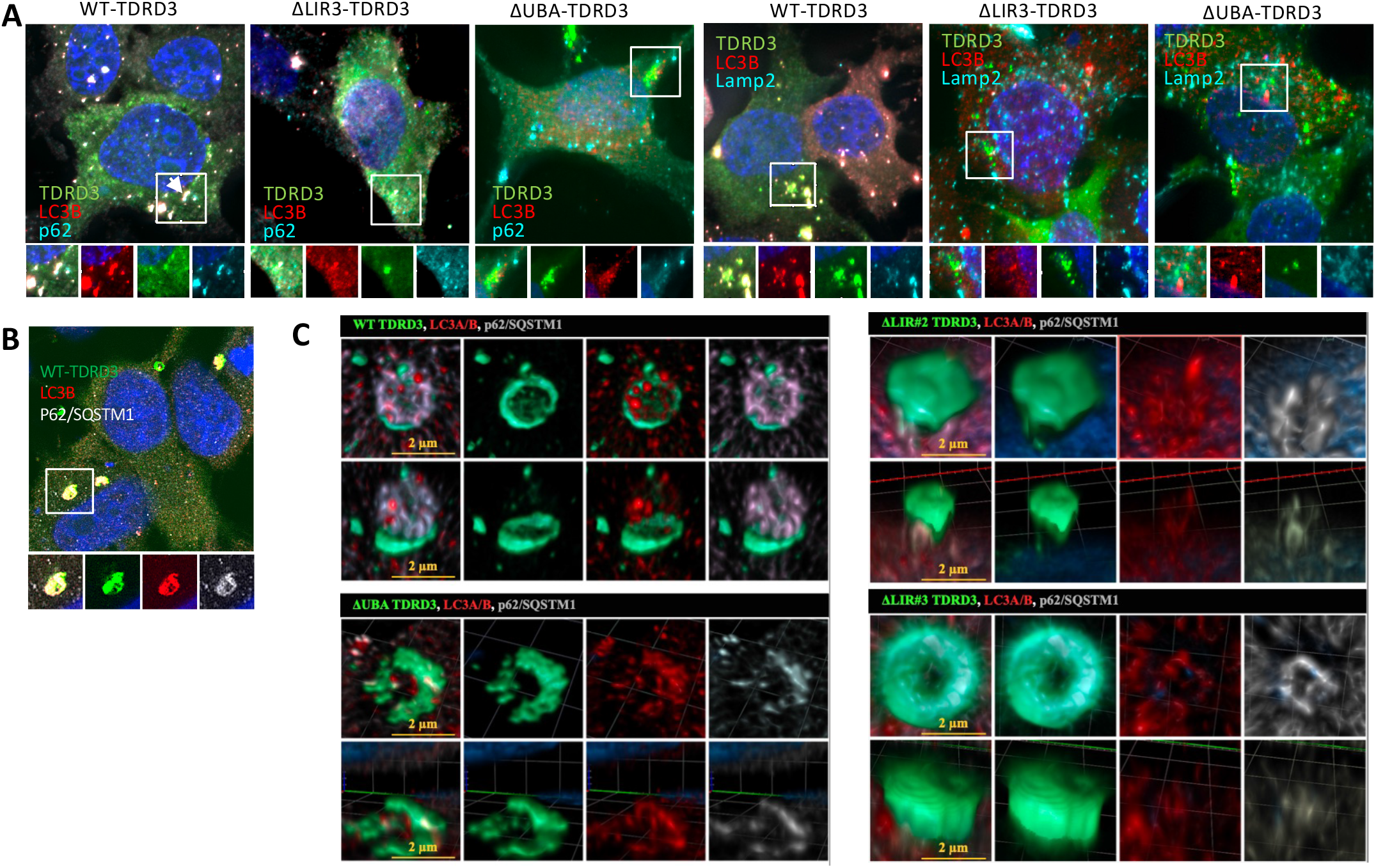
A. Colocalization of endogenous levels of TDRD3 variants with LC3B and p62/SQSTM1 during starvation. TDKO cells expressing low endogenous levels of TDRD3 variants were starved 4 hrs before fixation and processing for IFA microscopy by immunostaining for LC3B and p62/SQSTM1. Vignettes shown are taken from indicated areas in images. B,C. Starved cells processed by immunostaining for LC3B and p62/SQSTM1 were subjected to super-resolution microscopy.

When autophagosomes were imaged with super-resolution microscopy interesting structures were observed only during starvation suggesting complex interactions between TDRD3, LC3A/B and p62/SQSTM1 exist at these foci **(Fig. 4C)**. We observed that WT-TDRD3 participated in ring structures that were interfaced more closely with p62/SQSTM1 and LC3B. Examination of similar ring structures formed with mutant variants were all altered from this WT phenotype. ΔLIR2-TDRD3 did not form a similar ring, appearing thick and closed, with poor p62/SQSTM1 recruitment and only trace LC3B. ΔUBA-TDRD3 showed a heavier incomplete ring formation with stronger LC3A/B recruitment. The ring structure with LIR3-KO TDRD3 was large and heavy, with reduced p62/SQSTM1 recruitment. Taken together, these data suggest TDRD3 may participate in phagosome formation or maturation under limited conditions, and mutant variants form altered structures, consistent with displaying different defects in assembly or maturation.

### TDRD3 deletion variants affect stress granule morphology

TDRD3 is both recruited to SGs and over-expression induces SG condensates. We examined if expression of TDRD3 variants altered the morphology of SGs. Compared to normal SG which were mostly large and tightly condensed with G3BP1 and TDRD3, the expression of UBA-KO-TDRD3 exhibited more tightly condensed SGs **(Fig. 5A)**. The LIR1 and LIR2KO showed only modest changes from WT SG phenotypes. In contrast, LIR3 KO and LIR123KO developed SGs in small subset of cells only, possibly due to overexpression, and those were much less condensed or more dispersed than WT-TDRD3 SGs. Also, a substantial fraction of LIR3 KO TDRD3 failed to condense into significant SG condensates, remaining dispersed in the cytosol in numerous small TDRD3 foci that did not colocalize with G3BP**(Fig 5A, arrows)**. This suggests that the LIR3 motif may play a role in SG assembly. Alternatively, TDRD3 lacking this motif is readily stripped from SGs and directed to lysosomes. These smaller TDRD3 foci were also seen to lesser extent with LIR1 or LIR2KO TDRD3 expression. Thus, the various TDRD3 motifs differentially affected SG morphology and recruitment of TDRD3 to SGs with G3BP1. We have also identified the Tudor-domain-flanking LIR motif (LIR-3) as a key driver of TDRD3-SG recruitment during arsenite treatment.

**Fig. 5.**
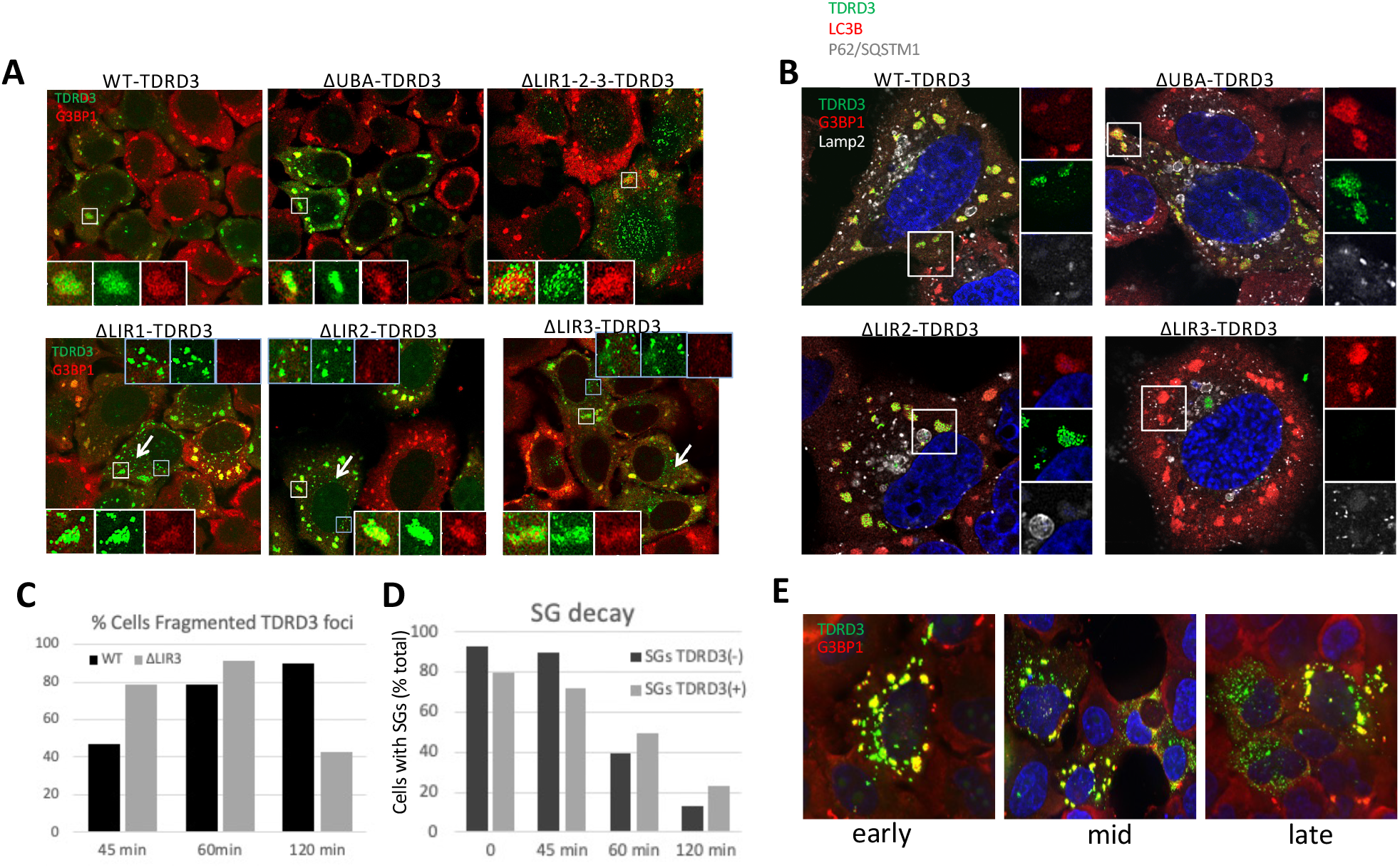
Colocalization of TDRD3 variants with LC3B and p62/SQSTM1 during arsenite stress and recovery from stress. A. IFA microscopy of TDRD3 variants expressed at endogenous levels in cells treated for one hr with arsenite, immunostained for G3BP1 (A) or LC3B and p62/SQSTM1 (B). Arrows indicate regions with numerous small TDRD3 foci that do not colocalize with G3BP1. C. The number of cells observed with similar small TDRD3 foci at indicated times post-Arsenite recovery. D. The number of cells containing G3BP1-stained SGs vs TDRD3-stained SGs (contain TDRD3 and G3BP1) at indicated times post-arsenite recovery. E. Examples of common phenotypes of TDRD3 condensates undergoing disassembly from SGs during 45, 60 or 120 min recovery.

### TDRD3 interfaces with autophagy machinery for SG maturation, stability, and dissolution

Since autophagy is proposed to eliminate SGs, we examined SGs formed during arsenite stress and for colocalization of autophagy markers. Overall we found no significant colocalization of LC3B with stress granules containing TDRD3 variants or not (DATA NOT SHOWN). Similarly, there was little significant colocalization of p62/SQSTM1 with SGs or during stress unless ring structures were found in certain cells (DATA NOT SHOWN, see Fig 6B)). In contrast, Lamp2 was found to be recruited to SGs, not dependent on TDRD3 since it is localized to SGs with G3BP1 but without TDRD3 **(FIG 5B** ) ΔLIR3-TDRD3 did not recruit to most SGs in many cells, but was often localized perinuclear in degradation centers with increased p62/SQSTM1 foci (**Fig. 5B**).

**Fig. 6.**
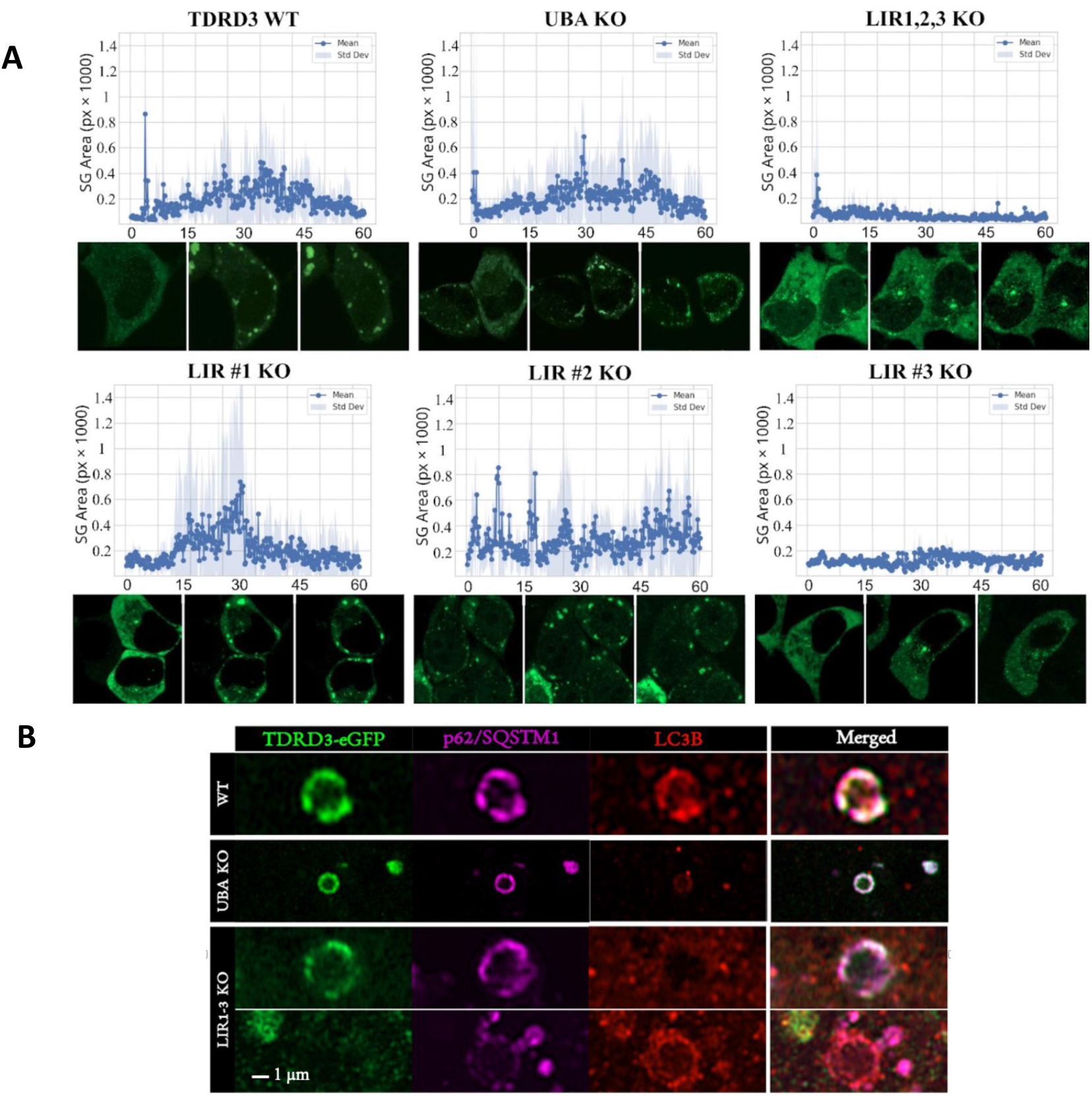
Kinetics of SG recruitment of TDRD3 deletion variants during arsenite induced SG assembly. Microscopy images were taken every 10 sec post-addition of arsenite for 60 sec and SG content determined by CellProfiler analysis. Images below graph are representatives from time 0, 30 min or 60 min. B. Super resolution microscopy of non-SG associated ring structures observed in some cells during Ars stress.

Since there was a defect in the morphology of SGs with mutant variant TDRD3 we performed analysis of the rate of SG disassembly**(Fig. 5D)**. Comparing cells not expressing TDRD3 to those expressing WT TDRD3, we found a slight reduction in the assembled SG, followed by sightly faster decay initially, then slower SG decay at 1 and 2 hrs. SG decay also produced smaller TDRD3 foci that did not colocalize with G3BP1. These were much more prevalent early in cells expressing LIR3-KO TDRD3 than those expressing WT TDRD3. A proposed sequential phenotype of steps of TDRD3/SG decay is shown, with TDRD3 fragmentation commonly appearing early, then smaller TDRD3 foci lacking G3BP1 as G3BP1 SGs diminish **(Fig. 7E)**.

**Fig. 7.**
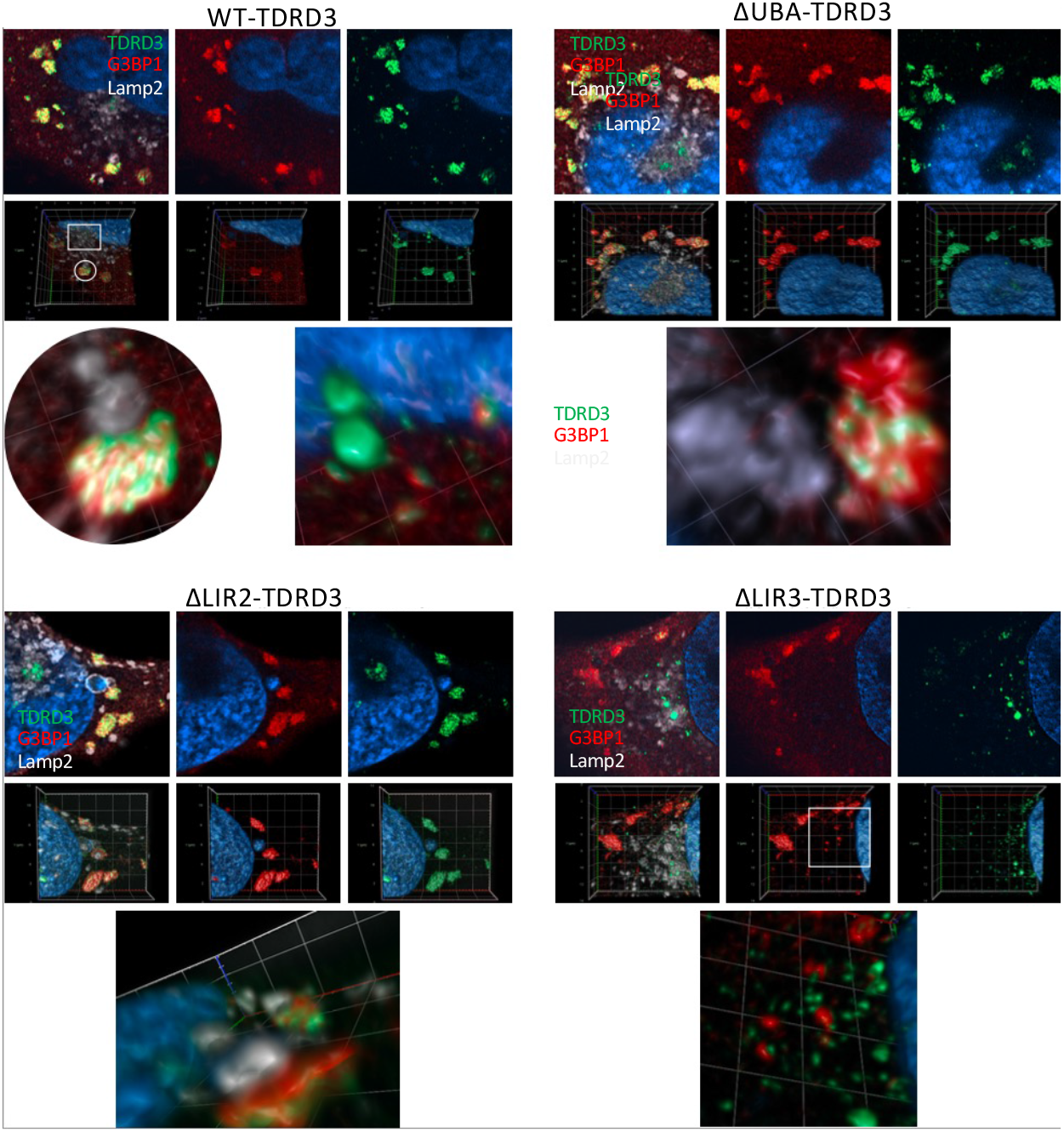
Super resolution microscopy of non-SG associated ring structures observed in some cells during Ars stress. Cells were expressing endogenous levels of the indicated TDRD3-eGFP variant. Top row for each indicated 2D micrograph, a 3D image of the same area is below that, followed by enlargements of indicated areas of 3D images.

### LIR3 dependent recruitment of TDRD3 to SG

SG resolution is proposed to involve autophagy, yet we find little colocalization of LC3B or p62/SQSTM1 with SGs. It is possible that TDRD3 functions in opposition to the selective autophagy of SGs, perhaps stabilizing SG condensates. Thus, we sought to determine whether or not TDRD3 interfaces with autophagy machinery to facilitate some aspect of SG formation or resolution kinetics. We selected for cells expressing inducible TDRD3 variants (WT, UBA deletion, 3x LIR deletion, and each individual LIR deletion) and then induced TDRD3 expression to near normal endogenous levels. We then stressed these cells with arsenite for an hour.

Using super resolution live imaging, we monitored the formation kinetics of SGs at 10 second increments over the course of 1 hour. We then analyzed each image to quantify each SG area (in pixels) over time as a measure of SG formation and maturation. We found that lacking either LIR-2 or LIR-3 (or all LIR motifs deleted) resulted in significant changes in SG maturation kinetics compared to WT. While the ΔUBA-TDRD3 variant produced SGs with similar kinetics to WT-TDRD3, appearing by 30 min, then showing a slight decline, LIR-2-TDRD3 was recruited to SGs a bit faster, and did not decline. (**Fig. 6**). LIR-3 was the most important recruitment factor for TDRD3 to SGs, as both variants of TDRD3 lacking this motif were very impaired in recruitment to SGs for the duration of the experiment.

Interestingly, we also found instances **(Fig. 6B)** where TDRD3 can interact with LC3B and p62/SQSTM1 to form ring like structures during arsenite stimulation similar to what was observed with EBSS induced starvation (see Fig 4B) although about half of the size (∼1 μm vs ∼2 μm during starvation). While WT- or ΔUBA-TDRD3 were found in these structures with both p62/SQSTM1 and LC3B, ΔLIR3-TDRD3 was found with p62/SQSTM1 alone or instances were found with p62/SQSTM1 together with LC3B but lacking ΔLIR3-TDRD3.

Finally super-resolution microscopy of Ars-treated cells revealed interesting structures. WT-TDRD3 was together with G3BP1 and Lamp2 vesicles apparently interfacing with TDRD3-enriched areas of the SG. In other parts of the cell TDRD3 was often perinuclear in two foci, possibly near microtubule organizing centers (mTOCs) **(Fig. 7)**. The ΔUBA-TDRD3 and ΔLIR2-TDRD3 could be seen interfacing with Lamp2 vesicles in a similar fashion, however the latter interface more strongly with G3BP1. In contrast, the ΔLIR3-TDRD3 variant was dispersed across the cytoplasm, not concentrated in SGs and did not interact with small G3BP1 foci.

## DISCUSSION

In this work we have discovered that TDRD3 is strikingly similar to known selective autophagy receptors, that it functions in autophagy by as of yet unknown mechanisms, and that reintroducing TDRD3 back into cells restores autophagy. We have confirmed that TDRD3’s LIR motifs as well as its UBA domain facilitate its interaction with the autophagy protein LC3B. Additionally, we have found that TDRD3 and p62/SQSTM1 are similarly degraded during autophagy but when lysosomal degradation is halted we observe an increase in both proteins suggesting that TDRD3, like p62/SQSTM1, likely function in selective autophagy. We also found that reintroducing TDRD3 into TDRD3 KO cells restores autophagy during starvation. In contrast to previous reports, we have found that it is TDRD3’s LIR motif (#3) that is largely responsible for its recruitment to SGs instead of its Tudor domain (Goulet 2008). Goulet et. al. actually deleted the full Tudor domain from TDRD3, which encompasses part of the LIR#3, thus it comes as no surprise that they attributed SG recruitment with this domain.

Interestingly, we found an association between TDRD3 and the size of LAMP2+ vesicles during oxidative stress, hinting that TDRD3 may be involved in the timely degradation of stress granules. During starvation, this phenomenon was not observed. However, G3BP1 appears to facilitate LAMP2+ vesicle sizes during starvation presenting the possibility that G3BP1 functions more in nutrient stress response. Indeed, we have found that G3BP1 also possesses an LIR within its unstructured region (amino acids 248-259) although we have not yet investigated its function. Surprisingly, via volumetric imaging we found that LAMP2+ vesicles interact very closely and may actually dock with TDRD3 in SGs.

TDRD3’s possible docking to LAMP2+ vesicles is an active area of current investigation as TDRD3 possesses 3 KFERQ-like sequences (retrieved from KFERQ finder VO.8 database^26^, one of which is a canonical motif in a disordered region that is neither requisitely activated by phosphorylation or acetylation and cannot be silenced by ubiquitin. This sequence is predicted to directly bind HSC70 which then docks to LAMP2 to unwind these KFERQ-containing proteins through LAMP2 and into lysosomes directly. This finding presents another possible mechanism by which TDRD3 may be associating with these LAMP2+ vesicles (Fig.7) for the purpose of possible engulfment of SGs into these vesicles via proximity or degrade SGs in a piecemeal fashion ultimately loosening SGs as SGs lacking TDRD3 appear “looser”(Fig. 5A) or even autophagocytosed SGs being led to these vesicles facilitating the proximity-based attachment of SNAREs on either vesicle in addition to charge negation imparted by RABs and their effectors^27^.

Intriguingly, we discovered that upon starvation TDRD3 forms large rings through which phagosomes appear to form. While further experimentation is required to determine the mechanistic underpinnings thereto, we have found that TDRD3 may associate with Sec16A (BioGRID ^28^), part of the Sec16 complex responsible for export of content from the ER at ER exit sites (ERES) (Reviewed in ^29^). An attractive idea is that TDRD3 may function in the maturation of autophagosomes. Additionally, TDRD3 has a putative association with dynactin (BioGRID ^28^), similarly to p62/SQSTM1 ^30^, thus it is also possible that TDRD3 may actually “walk” SGs toward the lysosome-rich perinuclear region. Surprisingly, we have discovered that TDRD3 accumulates at the MTOC suggesting that this may be how SGs traverse the microtubule network toward their eventual demise in lysosomes.

With regards to TDRD3’s potential role in the degradation of SGs, it is important to note that p62/SQSTM1 has been found to interact with SMN and symmetrically dimethylated arginine (SDMA) residues on FUS, marks deposited by PRMT5, resulting in the selective autophagy of SGs ^31^. TDRD3 is a known reader of asymmetric dimethylarginine (ADMA), and FUS enters SGs via PRMT1-deposited ADMA residues ^32^, thus it is possible that TDRD3’s potential recruitment of ADMA-FUS into SGs begins the process of SG resolution. This switching from ADMA to SDMA may also occur due to the presence of JMJD6 within SGs which we previously found to be involved in SG assembly via demethylation of G3BP1’s RGG domain^33^. TDRD3 has recently been shown to actually promote SG assembly in concert with PRMT1 ^8^, thus the idea that TDRD3 may truly function as a master regulator of SGs in concert with G3BP1 is attractive and will be investigated.

Previous work has shown SG clearance through autophagy involving CdC48/VCP using genetic approaches ^16,17^, that LC3B is an RNA-binding protein involved in RNA degradation^34^ and that G3BP1 ubiquitination mediates SG clearance ^35^. Here, we have shown that both TDRD3 and G3BP1 function in autophagy and most likely selective autophagy. We relate TDRD3’s function in this process to the presence of both its LIR motifs as well as its UBA domain. Interestingly, only LIRs 2 and 3 appear to function in TDRD3’s association with autophagy, while LIR #1 appears dispensable or we could not uncover a phenotype. This is likely due to the fact that this LIR (#1) is positioned within a structured domain, so accessibility may be completely blocked or function in a context-dependent manner not assessed here. The fact that the most relevant LIR motif, #3, is immediately adjacent to the Tudor domain of TDRD3 also presents the possibility that TDRD3 performs methyl-arginine binding or LC3 binding in a context dependent manner. The possibility that TDRD3 acts as a timer for stress response is further bolstered by this spatial association, similarly to the fact that LIR #2 is immediately downstream of the UBA domain. Interestingly, LIR#3 possesses a predicted phosphorylation site at its start which has been shown as an activating signal for other LIRs ^36^ and may play an important role in activating TDRD3 functions in autophagy.

## Materials and Methods

### Cells, transfections

Wild-type (WT) HeLa cells, G3BP1-knockout HeLa cells (G3BP1-KO) TDRD3-knockout Hela cells or G3BP1/TDRD3 double knockout HeLa cells were generated and used in this study. Where applicable, cells were transfected using Lipofectamine 3000 (Thermo Fisher Scientific) according to manufacturer’s instructions using reduced-serum Opti-MEM (Thermo Fisher Scientific) as the transfection media. For oxidative stress, sodium arsenite (Sigma) was added to culture media at 200 μM for 1 hour at 37°C.

### Plasmid constructs

pSI-TDRD3-eGFP, driven by an SV40 promoter was mutated using PCR (SuperFI II polymerase Thermo Fisher Scientific) to generate small deletions removing small ELMs from the TDRD3 sequence. Inducible constructs (pSII) were generated by cloning TetON-3G into MCS1 of pSI driven by an SV40 promoter. A secondary MCS (MCS2) was then generated downstream of the tet regulator. The Tet-Responsive TRE3G was cloned directly upstream of MCS2 to drive downstream expression of GFP or APEX2-tagged TDRD3 construct in a doxycycline-inducible manner producing an ectopically expressed all-in-one induction system. Variants of TDRD3 were created in the base plasmid pSI-TDRD3-eGFP by PCR-generated deletion of specific amino acid residues. TDRD3 Isoform 3 (744 amino acids) was used in all of this work. Knockout of the UBA domain deleted codons for amino acids 286-326, KO of LIR1 deleted amino acids 110-119, KO of LIR2 deleted for amino acids 364-374, KO of LIR3 deleted 645-651. All resulting plasmid constructs were verified by sequencing.

### Immunofluorescence(IFA)

Cells were grown on coverslips in 24-well plates. Cells were fixed in 2% paraformaldehyde and 0.2% glutaraldehyde diluted in PEM buffer (80 mM PIPES pH 6.8, 5 mM EGTA pH 7.0, 2 mM MgCl2) on ice for 45 minutes. Autofluorescence was quenched using sodium borohydrate (1mg/ml) diluted in PEM buffer twice for 5 minutes each followed by permeabilization for 10 minutes (PEM buffer supplemented with 0.5% Triton X-100) at RT. Cells were blocked for one hour at room temperature in 3% BSA in TBS supplemented with Triton X-100 (0.1%). Primary antibodies were diluted in blocking buffer. Cells were incubated with primary antibodies at 4°C overnight with rocking. After primary staining, coverslips were washed 5X with blocking buffer. Cells were incubated with either appropriate secondary antibodies conjugated to Alexa Fluor dyes (Invitrogen) at 1:500 concentration for 30 minutes at room temperature. Cells were then washed three times in TBS-TX100 followed by PEM buffer. Cells were then fixed again in 2% paraformaldehyde diluted in PEM buffer at RT for 15 minutes. Cells were then washed 3 times in PEM buffer for 5 minutes per wash, quenched twice in sodium borohydrate as previously performed, washed twice in PEM buffer and then washed once in TBS-T. Cells, where applicable, were then counterstained with Hoechst at 1:200 for 3 minutes then washed 3 times in TBS-T followed by TBS. Coverslips were then mounted using Prolong Glass non-hardening antifade mountant. Slides were then stored in the dark at 4°C until imaging. Imaging was performed on a Zeiss LSM980 with Airyscan 2 technology.

For live imaging, cells were transfected in 24-well plates as previous and then split into live imaging wells (ibidi, product 80826). Expression of inducible constructs was initiated by incubating cells in DMEM containing doxycycline (0.1 to 1 ug/ml) to sub-aggregation levels.

Opti-MEM I was used as a culture medium during imaging. Arsenite was added to Opti-MEM I and focal points were determined as cells expressing cytoplasmic GFP with no granules present (sub-aggregate induction levels). Imaging was performed using 4x multiplex low power feature capturing 1 image per 10 seconds over the course of 1 hour. Resulting images were then Airyscan processed to remove artifacts and enhance image signal.

### Western blot analysis and antibodies

Cell pellets from 1–8 x 10^6^ cells were lysed in RIPA buffer supplemented with HALT protease inhibitor cocktail (ThermoFisher) and phosphatase inhibitor (ThermoFisher) at 1X dilutions end-over-end tumbling at 4*C for 1 hour and then clarified by centrifugation (10 min., 10,000 RPM). Equivalent total protein concentrations (A_280_) were loaded per well and separated by SDS-polyacrylamide gel electrophoresis and transferred onto nitrocellulose membranes (Whatman). Blots were then blocked in Blotto (5% nonfat dry milk diluted in TBS-T) for 1 hour before incubation with primary antibodies overnight. Licor red and far red secondary antibodies were used at concentrations of 1:20,000. Western blot imaging was performed on a Licor CLx system. Antibodies used are listed below.

|  | <u>Host</u> | <u>Vendor</u> | <u>Catalog#</u> |
| --- | --- | --- | --- |
| LC3A/B | Rabbit | Cell Signaling Technology | 4108s |
| TDRD3 | Rabbit | Cell Signaling Technology | 5942s |
| p62 | Mouse | Proteintech | 66184-1-Ig |
| LAMP2 | Mouse | Invitrogen | 14-1078-82 |
| LC3B | Mouse | Invitrogen | MA5-37852 |
| G3BP1 | Rabbit | In-House | N/A |
| G3BP1 | Mouse | Santa Cruz | sc-365338 |

### RNA Extraction and qPCR

Cells were plated as described previously. Cells were treated with indicated treatments and then washed and lysed in-dish using Trizol reagent (Invitrogen) per manufacturer’s instructions. RNA was isolated and precipitated using 100% Isopropanol at -20*C for 20 minutes followed by centrifugation (12,000 g) for 15 minutes. RNA pellets were then washed and dried. Dried pellets were resuspended in RNAse-free water (Invitrogen). RNA was then treated with Turbo DNA-Free Kit (Invitrogen) per manufacturer’s instructions.DNA-free RNA was then quantified via spectrophotometry (Denovix DS-11). 1 μg of RNA was then converted to cDNA using random hexamers (Invitrogen) and Protoscript II (NEB) per manufacturer’s instructions. Resulting cDNA was then diluted to 12.5 ng/ul and 2 ul per well was used in subsequent qPCR. qPCR was performed using SYBR Green Master Mix (ThermoScientific #4309155) and detection was performed using the Applied Biosystems Quant Studio 3 standard qPCR protocol.

qPCR Primers used in this work:

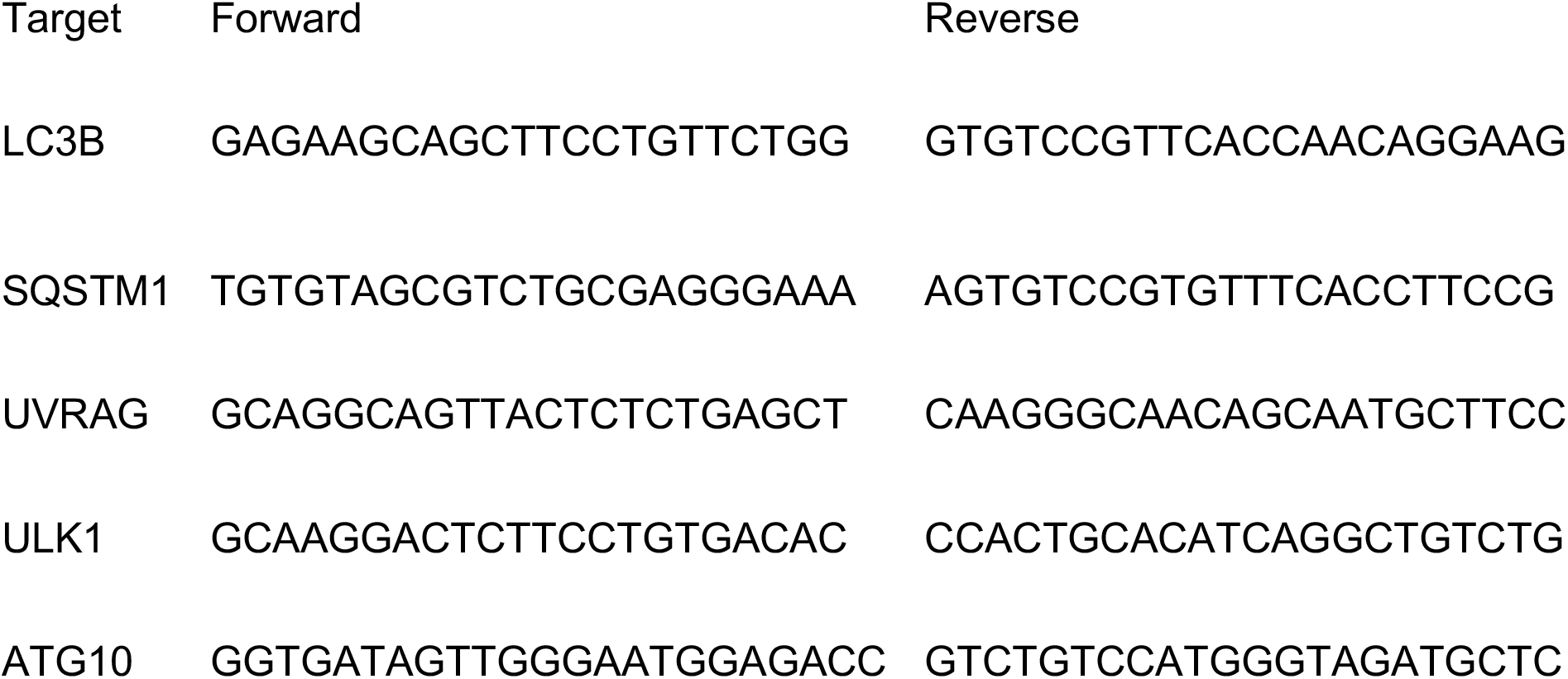

### Immunoprecipitation

TDRD3 KO cells were dual-transfected with GFP-RFP-tagged LC3B reporter (pBABE-puro mCherry-EGFP-LC3B, addgene #22418) and non-inducible TDRD3-APEX2 or indicated mutants thereof. Cells were incubated overnight and autophagy was stimulated using EBSS for 2 hours with no lysosomal degradation inhibitors. Cells were then lysed in CHAPS lysis buffer (0.5%, 300 mM NaCl in phosphate buffer, pH 7.4) to retain protein structures. Chromotek GFP trap was used according to manufacturer’s instructions to pull down LC3B with a GFP tag. Beads were washed with buffer containing 300mM NaCl to increase wash stringency. Samples were separated in 15-20% polyacrylamide gels and transfers were performed using 0.2 μm nitrocellulose membranes (ThermoScientific #88024) on ice at 250 mA for 75 minutes.

### Determination of LCII1/LC3II ratio

Immunoblots were processed to determine levels of LC3B-1 and II using specific antibodies (ref). Densitometry was performed using LICOR Image Studio 5. Statistical analyses was performed (Mann-Whitney-U-Test, one-tailed) in Python using SciPy and plotted using Seaborn. A minimum of biological triplicates were performed.

## Acknowledgements

This research was supported by National Institutes of Health grants AI150237 and S10 OD028480 (Zeiss LSM980-Airyscan 2 microscope).

---

**Supplemental Figure 1:**
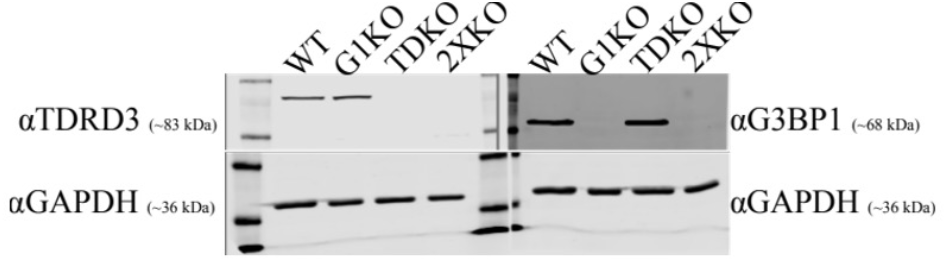
Western Blot Confirmation of TDRD3 KO in WT and G3BP1 KO HeLa Cells. TDRD3 KO was performed using All-In-One-CRISPR/Cas9_LacZ (Addgene #74293). Cells were selected with Neomycin for at least 2 weeks. Confirmatory western blotting is regularly performed.

**Supplemental Figure 2:**
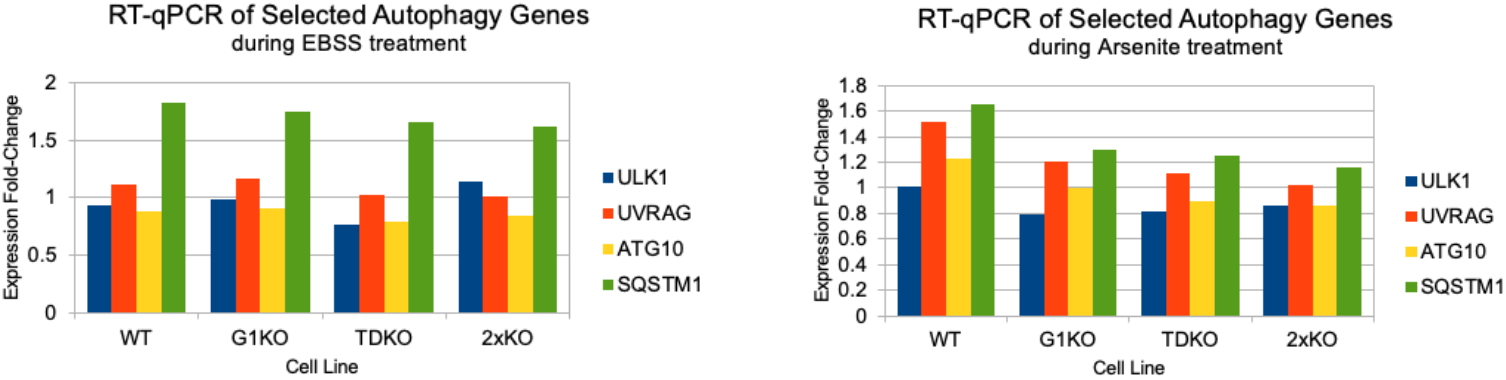
qRT-PCR analysis of mRNA levels of indicated autophagy related proteins in the cell lines indicated after treatment of the cell lines for starvation with EBSS for 4 hrs or oxidative stress by arsenite for 60 min.

